# Characterisation of the Renal Cortical Transcriptome Following Roux-en-Y Gastric Bypass Surgery in Experimental Diabetic Kidney Disease

**DOI:** 10.1101/2020.06.01.120980

**Authors:** Meera Nair, William P. Martin, Vadim Zhernovkov, Jessie A. Elliott, Naomi Fearon, Hans Eckhardt, Janet McCormack, Catherine Godson, Eoin Patrick. Brennan, Lars Fändriks, Neil G. Docherty, Carel W. le Roux

**Author notes:** **Correspondence to:** Dr. Neil Docherty, Diabetes Complications Research Centre, School of Medicine, Conway Institute of Biomolecular and Biomedical Science, University College Dublin, Belfield, Dublin 4, Ireland.

## Abstract

**Introduction:** Roux-en-Y Gastric Bypass Surgery (RYGB) reduces albuminuria and the long-term incidence of end-stage renal disease in patients with obesity and diabetes. Preclinical modelling in experimental diabetic kidney disease (DKD) demonstrates that improvements in glomerular structure likely underpin these findings.

**Research Design & Methods:** In adult male Zucker Diabetic Fatty (ZDF) rats, we profiled the effect of RYGB on weight and metabolic control as well biochemical, structural and ultrastructural indices of diabetic renal injury. Furthermore, we sequenced the renal cortical transcriptome in these rats and used bioinformatic pathway analyses to characterise the transcriptional alterations governing the renal reparative response to RYGB

**Results:** In parallel with improvements in weight and metabolic control, RYGB reduced albuminuria, glomerulomegaly, podocyte stress, and podocyte foot process effacement. Pathway analysis of RYGB-induced transcriptomic changes in the renal cortex highlighted correction of disease-associated alterations in fibrosis, inflammation and biological oxidation pathways. RYGB reversed disease-associated changes in the expression of TGF-β superfamily genes that strongly correlated with improvements in structural measures of glomerulopathy.

**Conclusions:** Improved glomerular structure in ZDF rats following RYGB is underpinned by pathway level changes, including interruption of the TGF-β driven pro-fibrotic programme. Our data provide an important layer of experimental support for clinical evidence demonstrating that RYGB arrests renal damage in patients with obesity and Type 2 Diabetes.

**KEY MESSAGES:**

1. What is already known about this subject?

- RYGB is an effective treatment for obesity and type 2 diabetes and longitudinal cohort studies have demonstrated it’s albuminuria lowering effect and evidence of longer term reno-protection.
- Studies in pre-clinical models of diabetic kidney disease have described favourable changes in measures of renal structure and ultrastructure following RYGB.
2. *What are the new findings?*

- The present study directly correlates structural and ultrastructural improvements in the ZDF rat kidney following RYGB with corrective shifts in the global renal transcriptome.
- Chronic renal remodelling responses in experimental DKD that are governed by TGF-β signalling are interrupted and reversed by RYGB.
3. *How might these results change the focus of research or clinical practice?*

- These data will support further interrogation of RYGB specific shifts in the renal transcriptome with a view to identifying tractable targets for treatment response biomarkers and bariatric mimetic based diet and pharmacotherapy based interventions.

## INTRODUCTION

In a 2017 analysis of the global burden of Chronic Kidney Disease (CKD), over 30·7% of associated disability-adjusted life years occurred in patients with background of diabetes mellitus, this being the largest contribution from any single cause. (1) Analysis of 2013-2016 U.S data from the National Health and Nutrition Examination Survey (NHANES) illustrate this point; with a microalbuminuria prevalence rate of over 40% in patients with an HbA1c >64mmol/mol (DCCT >8%). (2) Approximately 25% of patients in this category have an estimated glomerular filtration rate (eGFR) of less than or equal to 60ml/min/1.73m^2^. (2) Longitudinal analysis of NHANES III data reveals that patients developing progressive albuminuria and eGFR decline have an estimated 10-year all-cause mortality incidence rate of 47%, 4-times higher than the rate observed in patients with diabetes and no evidence of renal impairment. (3)

The United Kingdom Prospective Diabetes Study (UKPDS) demonstrated the independent and additive impact of targeting hyperglycaemia and hypertension on microvascular outcomes of type 2 diabetes mellitus (T2DM) at medium-long term follow-up. (4) Multimodal lifestyle and pharmacology based protocols such as those deployed in the STENO-2 trial emphasise the benefits of comprehensive risk-factor control. (5) Recently developed glucose-lowering therapies, including glucagon-like peptide 1 receptor agonists (GLP1RAs) and sodium glucose co-transporter 2 inhibitors (SGLT2is), improve cardiovascular and renal outcomes in people with diabetic kidney disease (DKD). (6–9) Nevertheless, DKD remains a progressive disease.

Bariatric surgery is an effective intervention with long-term weight-loss and glucose-lowering efficacy in patients with obesity and type 2 diabetes. (10–13).

A substantial evidence base now exists in the literature pointing to improved long-term cardiovascular and renal outcomes following bariatric surgery. (14). A recent metanalysis including data from 17,532 patients drawn from 10 high quality clinical studies reported that bariatric/metabolic surgery was associated with significant improvements in DKD versus medical treatment (odds ratio 15·41, 95% CI 1·28 to 185·46; P =0·03). (15) Much of the benefit observed was premised upon reductions in urinary albumin excretion, supporting previous focussed meta-analysis of the beneficial impact of bariatric/metabolic surgery on this parameter in patients with DKD. (16) Two further key reports have emerged in 2020, which add significant weight to the clinical evidence base. Firstly, in a balanced comparison of the impact of metabolic surgery (Teen-LABS study NCT00474318) versus active lifestyle and medical therapy (TODAY study NCT00081328) on DKD in adolescents with obesity and T2D, marked increases in odds ratios for hyperfiltration (OR 17.2, 95% CI 2.6–114.5; P=0.03) and albuminuria (OR 27.3, 95% CI 5.2–146.2: P<0.0001) were noted over 5 year follow-up in the TODAY participants. (17) This reflected opposing trajectories in these parameters over follow-up. Secondly, we have recently collaborated in the *Clinical Study on Metabolic Surgery Compared to the Best Clinical Treatment in Patients with Type 2 Diabetes Mellitus* (*MOMS*) RCT (NCT01821508). (18). This study demonstrated that at 2 year follow-up in patients with T2DM, Class I obesity (BMI 30-35kg/m^2^) and mild-moderate DKD (albumin-to-creatinine ratio [uACR] 3-30mg/mmol and eGFR >30mL/min/1.73m^2^) microalbuminuria remission occurred in 82% (95% CI=72-93%) of patients after RYGB versus 54.6% (95% CI=39-70%) of patients after best medical treatment (P=0.006). (18)

The potent glucose lowering effects of Roux-en-Y gastric bypass surgery (RYGB) have been demonstrated to be complimented by improvements in blood pressure which may be mechanistically linked in part to a unique natriuretic effect of the procedure. (19–20) We have previously reported that RYGB effectively reduces albuminuria in the Zucker Diabetic Fatty (ZDF) rat in conjunction with evidence of improvements in glomerular histology and ultrastructure. (21, 22)

Herein we present an in-depth analysis of global changes in the renal cortical transcriptomic occurring in response to RYGB in ZDF rats. We report that improvements in renal injury following RYGB in the ZDF are underpinned by significant pathway levels shifts in the renal cortical transcriptome that correlate with structural improvements, cross-reference to human DKD and highlight that RYGB interrupts the pro-inflammatory and TGF-β driven fibrotic programme characteristic of progressive CKD.

## METHODS

### Animal Studies

Experiments were undertaken under governmental project license (Health Products Regulatory Authority– AE18692-P084). ZDF (fa/fa) and fa/+ rats (Charles River Laboratories (France and U.K.)) were purchased at 6 weeks of age and maintained on Purina 5008 chow *ad libitum*. Experimental groups consisted of fa/+ rats (n=6), SHAM-operated fa/fa rats (n=8), and RYGB-operated fa/fa rats (n=9). Surgery was performed at 12 weeks of age and animals were humanely killed at the end of the eighth post-operative week. Body weight and glycaemia (ACCU-CHEK^®^-Roche) were assessed weekly.

### Surgical Protocols

Insulin degludec (Novo Nordisk, Denmark) was used for 7-days prior to surgery to achieve morning glucose readings of <12 mmol/L. Under isoflurane anaesthesia, a midline laparotomy and mobilisation of the gut was performed as SHAM surgery. According to standard protocol (21, 22), in RYGB-operated rats the proximal jejunum was transected 12cm from the pylorus and a side-to-side jejuno-jejunal anastomosis created leaving a common channel of 30cm. A small gastric pouch was created and end-to-end anastomosed to the alimentary limb.

### Biochemical Analyses

Albumin-Creatinine Ratio (ACR)- Urinary albumin and creatinine were assessed in urine samples collected over 16 hours in metabolic cage. Metabolic cage studies were undertaken the week prior to surgery and 8 weeks later. Analytes were measured using an autoanalyzer (Roche/Hitachi Cobas c502 Modular analyser) and resulting values used to derive the ACR (mg/g). Colorimetric assay was used to measure mg/dL concentrations of plasma triglyceride (Cayman Chemical 10010303) and total cholesterol (WAKO 294-65801). Urinary osteopontin concentrations (ng/ml) were assessed by ELISA (R&D DOST00), normalised by urinary creatinine and reported as ng/mg.

### Glucose Tolerance Test

RYGB operated ZDF rats and fa/+ animals underwent glucose tolerance tests at 19 weeks of age, 8 weeks after the ZDF rats had underwent RYGB. Animals were fasted for 16 hours before receiving a bolus of glucose (1g/kg), intra-peritoneally or orally. Blood glucose measurements were taken at time 0, 15, 30, 45, 60, 90 and 120 minutes following bolus administration. Area under the curve was calculated using the trapezoidal rule.

### Histological & Immunohistochemical Analyses

Immunohistochemical studies were carried out using 4μm thick sections of formalin fixed paraffin embedded kidney or pancreas. Picrosirius Red staining of kidneys was carried out on 10μm thick sections using a kit (Polysciences 24901). Immunohistochemistry of Wilms’ Tumour (WT-1), (antibody C-19, sc-192 Santa Cruz Biotechnology, 1:250 dilution) and desmin (Antibody Clone Dako.Cytomation 1:100 dilution) and insulin (antibody ab181547 Abcam, 1:64,000 dilution) were carried out using a Dako Autostainer Link 48 system. Signal amplification was achieved using the horseradish peroxidase-based FLEX system with signal development using 3,3’-Diaminobenzidine (DAB)+ substrate. IgG1 isotype controls and no antibody controls were used to confirm specificity of staining.

Glomerular morphometry was performed using Image J 1.48v software analysis of scanned 2Ox images of WT-1 stained sections (Hamamatsu Photonics, Digital Slide Scanner). Thirty glomeruli per sample were analysed. Glomerular volume (μM^3^) was estimated using the Weibel and Gomez formula. (23) Quantification of desmin staining was carried out in 25 individual glomeruli per sample. Desmin and insulin staining was quantified using an algorithm with macros specific to DAB staining derived from the Aperio Cytoplasmic V2 Algorithm (Leica).

### Transmission Electron Microscopy (TEM)

Glutaraldehyde fixed and 1% osmium tetroxide post-fixed renal tissue was dehydrated and infiltrated with EPON™ Epoxy Resin. Ultra-thin sections were prepared and imaged by transmission electron microscopy (Technai 12). Glomerular basement membrane (GBM) thickness was measured according to the Haas method at 22500x (24). Twenty GBM thickness measurements were recorded per specimen, from a minimum of five separate capillary loops and representative of at least three separate glomeruli per sample. Podocyte Foot Process Frequency (PFPF) was estimated at 9900x by determining the number of podocyte foot processes per unit length (8μM) of peripheral GBM. Podocyte Foot Process diameter (PFPD) was also measured as a reciprocal. Six measurements were recorded per sample.

### Transcriptomic and Bioinformatic Analyses

RNA isolation was carried out using RNeasy mini kits (Qiagen). RNA concentration and quality were measured using NanoDrop 2000 (Thermo Fischer) and bioanalyzer analyses (Agilent 2100) respectively. An RNA integrity number (RIN) of >7 was set as the quality threshold for subsequent RNA sequencing. RNA Library preparation was carried out using the TruSeq^®^ Stranded mRNA NeoPrep™ Kit (Illumina NP-202-1001). Libraries were pooled and diluted to 1.8pM then sequenced (NextSeq-lllumina FC-404-2002). Raw data processing and differential expression analysis was carried out with the support of The Bioinformatics Core Facility, University of Gothenburg, Sweden followed by pathway enrichment analyses using the R programming language (Supplementary File S1). Sequencing data files have been deposited in GEO (GSE117380)

### qPCR and Western Blotting

Validation of selected genes was conducted at the mRNA level by TaqMan quantitative reverse transcriptase PCR (Thermo-Fischer). Amplification was performed on QuantStudio 7 Flex^®^ (Applied Biosystems). Beta-actin was used as an endogenous control for normalisation of target genes. Results were analysed based on ΔΔCt quantitation, method of analysis. For Western blot analysis, normalized protein extract was resolved by SDS-PAGE. Proteins were then transferred onto PVDF membranes (Millipore), blocked with TBS-T supplemented with 5% (w/v) non-fat dried milk or BSA and then incubated with the following antibodies at 4°C overnight: Alpha-tubulin (1:10,000 in 5% milk; Abcam Ab4074), Fibronectin (1:5,000 in 5% milk; Abcam Ab2413), and Vimentin (1:1,000 in 5% BSA; Cell Signalling Rabbit mAb #5741). Membranes were subsequently incubated with horseradish peroxidase-linked secondary antibodies for 1 hour at room temperature (Cell Signalling) and developed using chemiluminescence reagents TM (WesternBright^™^).

### Descriptive and Inferential Statistics

Analyses were conducted using GraphPad Prism V.6 and IBM SPSS V.24. Individual data points for each group are presented in boxplots with annotation of the group median and inter-quartile range (IQR). Longitudinal repeated measures data was assessed by Wilcoxon signed-rank test. Cross-sectional data was assessed by Kruskal-Wallis testing, correcting for multiple comparisons using Dunn’s test. Statistical significance was set at p<0.05.

## RESULTS

### Weight Loss and Improvements in Metabolic Control Following RYGB (Figure 1)

Despite maintaining a higher bodyweight for the majority of the study period, by study close, there was no significant difference in body weights between SHAM-operated ZDF rats and fa/+ lean controls 363.5g (332.5-383.3) vs. 386.5g (374.8-413.3); p=0.22, Figure 1A). Bodyweight in the RYGB group was significantly lower than in the SHAM group 327g (316-340) vs. 386.5g (374.8-413.3); p<0.001) and not significantly different from weights in the fa/+ group (p=0.31, Figure 1A). Over the course of the intervention period, median AUC calculated from the sum of morning plasma glucose readings in the SHAM group was 4-fold greater than that recorded in the fa/+ group (283.0 (237.8-338.4) vs. 71.7 (69.36-72.81); p<0.001, Figure 1B). Median AUC plasma glucose in the RYGB group, although elevated relative to the fa/+ group, was significantly lower than in the SHAM group (164.3 (137.7-184.4) vs. 283.0 (237.8-338.4) p=0.03). Morphological examination of the pancreatic islets was carried out alongside immunohistochemical detection of pancreatic insulin expression (representative images in Figure 1C). Islets in in the fa/+ group were of a round, uniform morphology. In contrast, islets in the SHAM group were less uniformly round, and often had a disrupted appearance. Islets in the RYGB group were more uniform. Insulin staining in the islets of SHAM operated animals demonstrated a trend to be lower relative to the fa/+ group with the median number of insulin positive cells/mm^2^ in the SHAM group being 2231 (1413-2634) and in the fa/+ group 3304 (3135-4354) (p=0.053). The median number of insulin positive cells per mm in the RYGB group was 4480 [IQR 3548-4833] and hence insulin staining was significantly increased in the RYGB islets relative to the SHAM group (p=0.008). Although there was no difference in insulin staining between the RYGB and fa/+ groups (p>0.999) and morning glucose levels were largely normalised after RYGB (Figure 1B), evidence of ongoing beta-cell dysfunction in RYGB operated animals relative to fa/+ was observed by conducting oral and intra-peritoneal glucose tolerance tests at 8-week follow-up (Figure 1D).

**Figure 1-.**
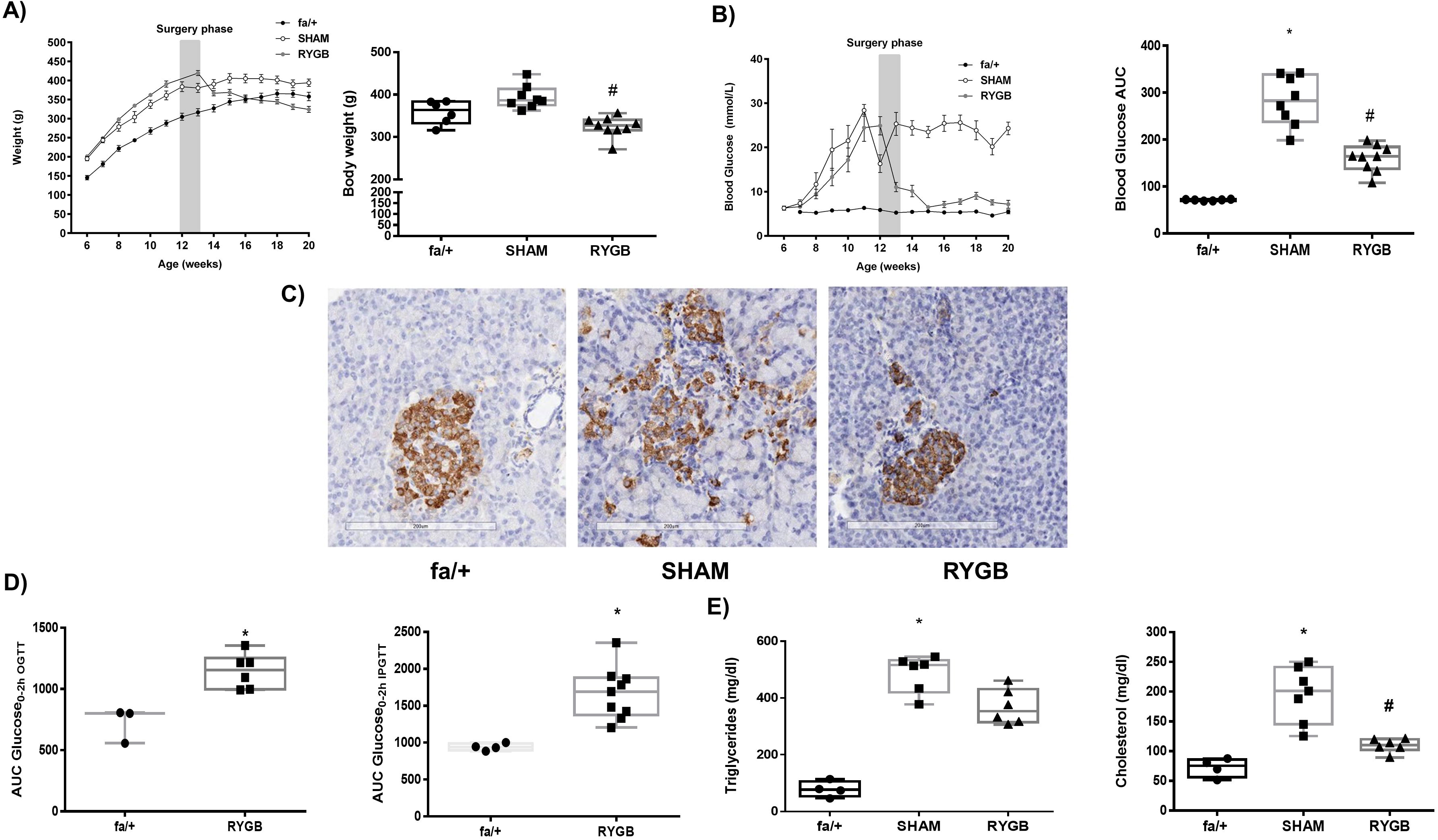
RYGB Reduces Bodyweight and Improves Metabolic Control. Trajectories of A) Body weight (g) and B) morning plasma glucose concentrations (mmol/L) before and after surgery and final values at post-operative week 8. C) Representative images of insulin immunohistochemistry based assessment of pancreatic islets at post-operative week 8 showing improved morphology and insulin positivity post-RYGB. D) Oral and intraperitoneal glucose tolerance testing in fa/+ rats and ZDF fa/fa rats at 8 weeks post-RYGB. Glucose tolerance tests were not performed in SHAM operated ZDF rats due to predictable futility given severe fasting hyperglycaemia which exceeded predicted maximal excursions in fa/+ rats or ZDF RYGB operated animals E) Plasma triglyceride (mg/dl) and total cholesterol (mg/dl) concentrations at post-operative week 8.) *p<0.05 vs fa/+; #p<0.05 vs. SHAM. SHAM-sham surgery (laparotomy); RYGB-Roux-en-Y Gastric Bypass, OGTT-oral glucose tolerance test, and ipGTT-intraperitoneal glucose tolerance test).

Median plasma triglyceride concentration was elevated in the SHAM group relative to the fa/+ group 515.6mg/dL (419.5-8-532) vs. 76.8mg/dL (53.2-105.7);p=0.01, Figure 1E). Triglyceride levels were not lowered relative to SHAM in the RYGB group. Median plasma cholesterol was 2.7-fold higher in the SHAM group relative to the fa/+ group (201.1mg/dL (145.2-241.3) vs. 75.3mg/dl (56.1=87.5); p<0.001, Figure 1E). Plasma cholesterol levels were reduced in the RYGB group relative to SHAM group (110.3 (101.9-121.1) vs. 201.1mg/dL (145.2-241.3); p=0.03 Figure 1E).

### Improvements in Albuminuria, Glomerular Structure and Ultrastructure Following RYGB (Figure 2)

Longitudinal paired analysis of reduced urinary albumin-to-creatinine ratios (uACRs) (Figure 2B) after RYGB showed reductions at 8 weeks post-surgery relative to baseline pre-operative values (17.7μg/mg (9.5-55.7) vs. 159.4μg/mg (10-468.8); p=0.010). Median glomerular volume (Figure 2C) was elevated in the SHAM group relative to the fa/+ group (1.2×10^6^μM^3^ (1.1×10^6^-1.5×10^6^) v’s 8.1×10^5^ μm^3^ (7.61×10^5^-8.6×10^5^); p=0.005. Median glomerular volume in the RYGB group was reduced by 33% relative to the SHAM group (p=0.020) and was not different to the fa/+ group (p>0.999).

**Figure 2-.**
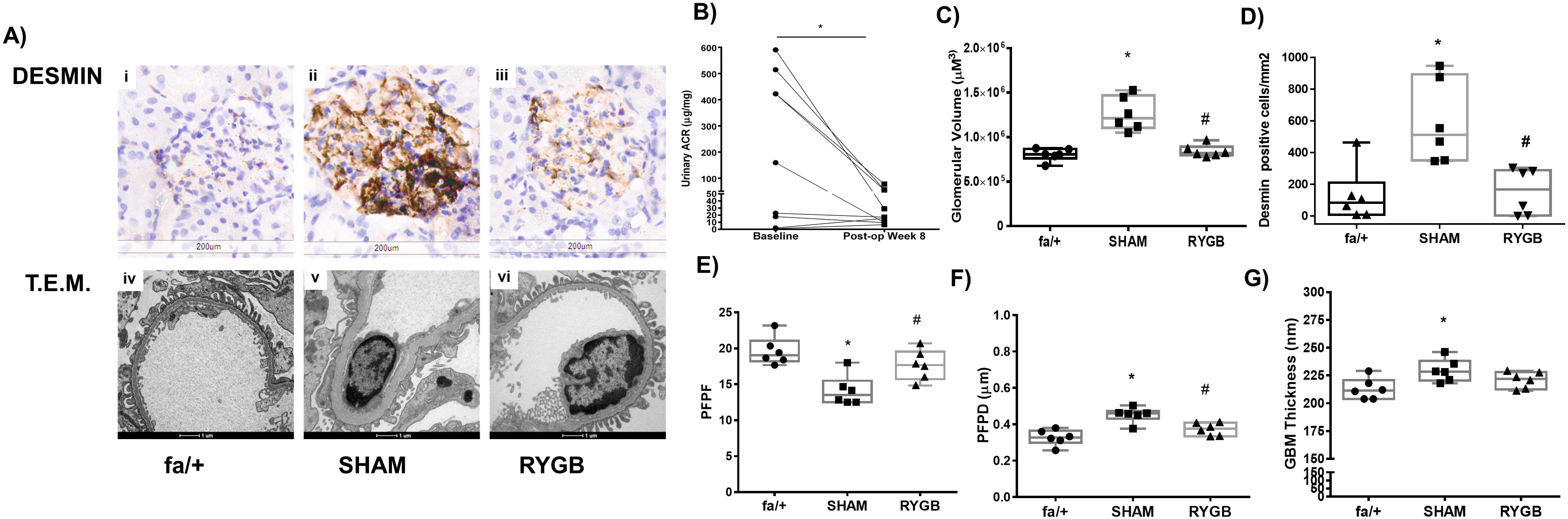
Impact of RYGB on Biochemical, Structural and Ultrastructural Indices of Diabetic Renal Injury. A) Representative images of glomerular desmin expression (immunohistochemistry 20x, scale bar 200μM) and glomerular ultrastructure (TEM, 9900x, scale bar 1μM) in fa/+, SHAM and RYGB rats. B) Urinary ACR (μg/mg) during the week before surgery and at post-operative week 8. C) Glomerular volume was assessed in 30 glomeruli per animal and the median glomerular volume (μM^3^) calculated. D) Glomerular desmin positivity (cells/mm^2^) was assessed in 25 glomeruli per animal. E) PFPF, F) PFPD and G) GBM thickness were assessed using T.E.M. For B p<0.05 vs.baseline. For C through H *p<0.05 vs fa/+; #p<0.05 vs. SHAM. SHAM-sham surgery (laparotomy); RYGB-Roux-en-Y Gastric Bypass, TEM-Transmission Electron Microscopy, ACR-Albumin Creatinine Ratio, PFPF-Podocyte Foot Process Frequency, PFPD-Podocyte Foot Process Diameter, GBM-Glomerular Basement Membrane).

Glomerular desmin staining (Figure 2D) was increased in distribution and intensity in the SHAM group compared to fa/+ and RYGB. Staining was localized within the glomeruli, in the mesangium, and at the periphery of the glomerular tuft. Median glomerular desmin positivity (Figure 2D) was elevated 6-fold in SHAM relative to the fa/+ group (512.1cells/mm^2^ (350.8-893.9) vs. 84.4 cells/mm^2^ (8.3-211.5); p=0.021). Glomerular desmin positivity was significantly lower in RYGB relative to SHAM (168.6cells/mm^2^ (4.2-289.2) vs 512.1cells/mm^2^ (350.8-893.9) ‘p=0.014). There was no difference in desmin staining between the fa/+ and RYGB group (p>0.999).

Podocyte foot process frequency (PFPF, Figure 2E) was reduced by 30% in the SHAM group relative to the fa/+ group (13.5 (12.5-15.5) vs. 19.0 (18.2-21.0); p=0.002). Median PFPF in the RYGB group was higher than in the SHAM groups (17. (15.71-19.54) vs 13.5 (12.5-15.5); p=0.026) Accordingly median podocyte foot process diameter (PFPD, Figure 2F) was 40% higher in the SHAM group relative to the fa/+ group (0.46μM (0.43-0.47) vs. 0.33μM (0.30-0.36); 0.33±0.04μM; p=0.002, with measures in the RYGB group significantly lower than in SHAM (p=0.015). Increases in glomerular basement membrane (GBM) thickness (Figure 2G) were observed in the SHAM group relative to the fa/+ group (228.4nm (220.3-238.2) vs. 211.3nm (203.9-220.8); p=0.044). Median GBM thickness in the RYGB group was intermediate to those of fa/+ and SHAM at 220.0nm (212.5-227.9).

### RYGB Corrects Disease-Associated Pathway Level Changes in The Renal Cortical Transcriptome (Figure 3)

Figure 3A shows dimension reduction of variance in the sequencing data along the first 2 principal components. Separate clustering of fa/+ and RYGB derived samples is evident against a background of relatively increased variance in the SHAM group. A total of 379 genes were identified as being differentially expressed in the ZDF rat kidney versus the fa/+ (212 downregulated, 167 upregulated, absolute fold-change ≥1.3 and adjusted p-value<0.05-Supplementary Table S2), corresponding to a change in 2.1% (379/18,423) of the common renal transcriptome. Genes showing marked changes in expression in the kidneys of diabetic animals included cardinal indicators of tubular injury and inflammatory and fibrotic change such as such as *Umod* (uromodulin), *Havcr1* (Kidney-Injury Molecule 1), *Vim1* (vimentin), *Fn1* (fibronectin), *Spp1* (osteopontin) and *IL24* (interleukin-24)

**Figure 3.**
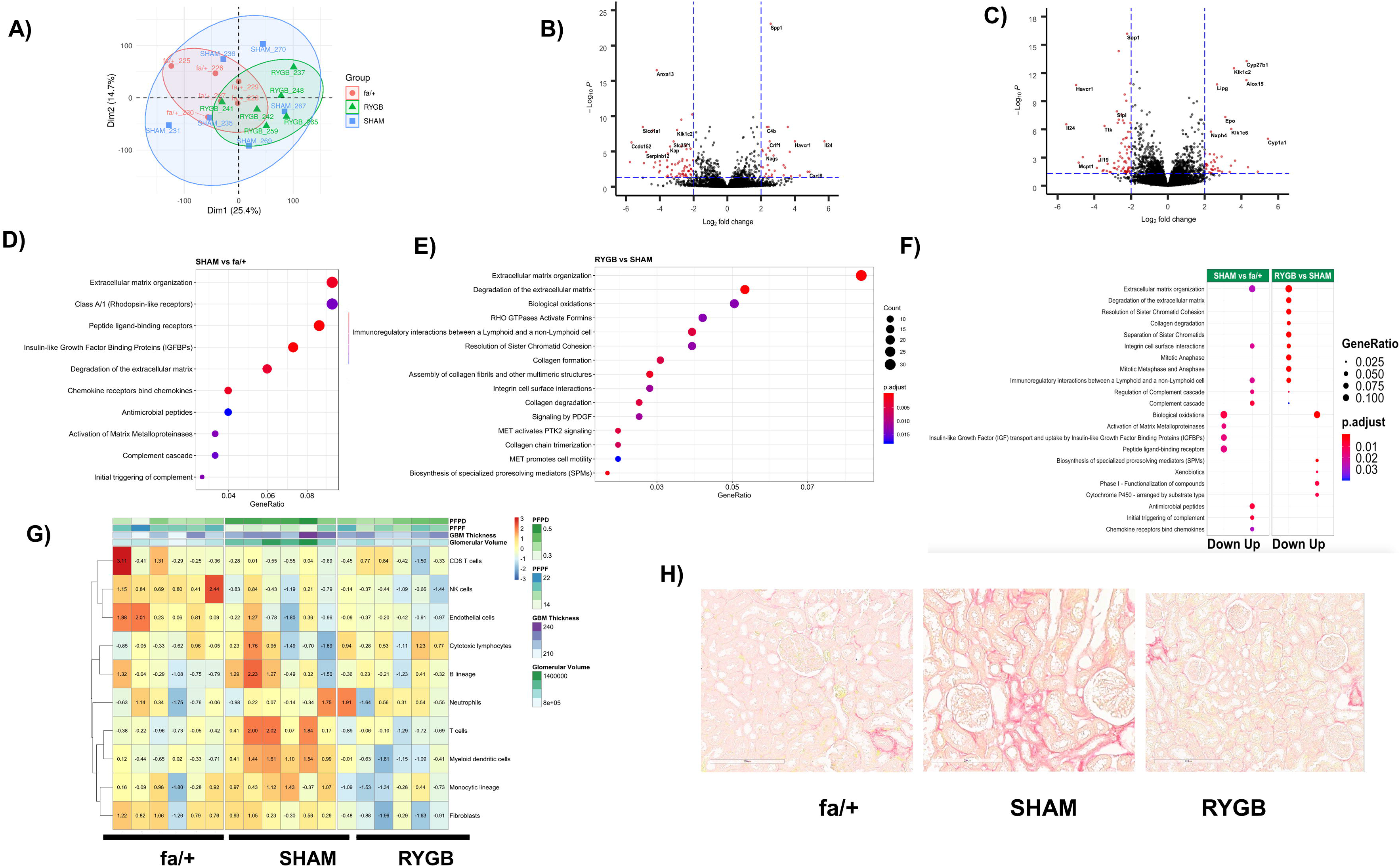
Reductions in Fibrotic Pathway Activation Dominate the Transcriptomic Response to RYGB. A) Principal component analysis based summarisation of the renal cortical transcriptome in fa/+, SHAM and RYGB operated rats. Volcano plot visualisation of differential expression analysis for the SHAM versus fa/+ comparison (B) and RYGB versus SHAM comparison (C). Reactome based analysis of pathway level changes in the SHAM versus fa/+ (D) and RYGB versus SHAM comparisons (E). F) Alignment of pathway level changes between SHAM versus fa/+ and RYGB versus SHAM comparisons indicating directionality of change. A fold-cut off change of 1.3 was applied in pathway analyses. In each graphic, adjusted p-value and gene ratios for each significantly altered pathway are represented as a function of colour and size of dots respectively. G) Heat-map visualisation of MCP Counter based estimates of the abundance of 8 immune and 2 stromal cell populations. Data is overlaid with heat-map annotation of quantitative measures of glomerular structure and ultrastructure for each sample included in the transcriptomic analysis. Colour coding indicates per sample relative abundance as a function of centered and scaled row specific data. Blue indicates low relative abundance and orange-red increasing relative abundance H) Representative images of Picrosirius-Red staining for total collagen in renal cortex. (SHAM-sham surgery (laparotomy); RYGB-Roux-en-Y Gastric Bypass).

The Comparison of the transcriptome of SHAM-and RYGB-operated ZDF rats 8 weeks after surgery identifies a total of 942 genes as being identified differentially expressed (529 downregulated, 413 upregulated, absolute fold-change ≥1.3 and adjusted p-value<0.05 Supplementary Table S3). This corresponds to 5.1% (942/18,423) of the common renal transcriptome. Reciprocal counter-regulation between SHAM versus fa/+ and RYGB versus SHAM of the expression of recognised disease associated genes such as those mentioned above is evidenced by pairwise comparison of Volcano plots (Figures 3B and 3C). RYGB was also associated with augmented expression of several genes that likely reflect adaptive responses to sub-optimal micronutrient assimilation following RYGB, including *Epo* and *Cyp27b1*, indicative of impaired iron and vitamin D homeostasis, respectively.

Disease-associated enrichment of multiple pathways in the Reactome database was detected (Figure 3D), with notable emergence of pathways linked to the innate immune activation and extracellular matrix remodelling. Enrichment of changes in extracellular remodelling, inflammation, cell cycle and biological oxidation related pathways were highly prominent in the RYGB versus SHAM comparison (Figure 3E). Reciprocal changes at pathway level between SHAM versus fa/+ and RYGB versus SHAM comparisons were evident (Figure 3F), suggesting arrest of progressive inflammation and fibrosis as well as restoration of tubular biotransformation function following RYGB. Interestingly, enrichment in cell cycle-related pathways after RYGB appeared as a discrete phenomenon not directly reflecting a correction away from the disease-associated profile of SHAM-operated kidneys. Congruent with pathway analysis, animals that underwent RYGB surgery were predicted to have a relative reduction in immune cells and fibroblasts versus SHAM-operated animals using MCP-counter (Figure 3G). Picrosirius Red (Figure 3H) demonstrates that SHAM-operated animals showed characteristic signs of early fibrosis including capsular fibrosis and patchy collagen deposition in the tubulointerstitium at sites of strong inflammatory infiltration adjacent to damaged tubules. Such foci were not uniformly observed in the kidneys of animal that underwent RYGB.

### RYGB Normalises the Expression of Genes that Correlate with Markers of Renal Injury and Cross-reference to the Human DKD Transcriptome (Figure 3)

We subsequently focussed on the extent to which RYGB corrected characteristic disease-associated gene expression patterns in the SHAM-operated ZDF rat. The Venn Diagram in Figure 4A depicts 144 differentially expressed genes common to SHAM versus fa/+ and RYGB versus SHAM comparisons. The directionality of change following RYGB was opposite to that occurring between health and disease in all but one case (RANTES), highlighting that 38% of the major corrective impact of RYGB on the renal cortical transcriptome (Figure 4B). We cross-referenced the list of 144 transcripts to the the Woroniecka human DKD glomerular microarray dataset (10), identifying 22 transcripts which were significantly differentially expressed in both human DKD and ZDF SHAM rats and which were also counter-regulated following RYGB (Figure 4C). From within this subset of genes translating to human disease, relative levels of mRNA and protein expression for vimentin, fibronectin and Spp1/osteopontin were assessed as a validation of the RNA sequencing analysis (Figure 4D and 4E). When normalised gene expression counts for transcripts within the group of 22 genes translating to human disease were correlated with markers of glomerular injury (Figure 4F), strong correlations with disease status were noted thereby associating these specific transcriptional shifts following RYGB with quantitative changes in glomerular structure and ultrastructure.

**Figure 4-.**
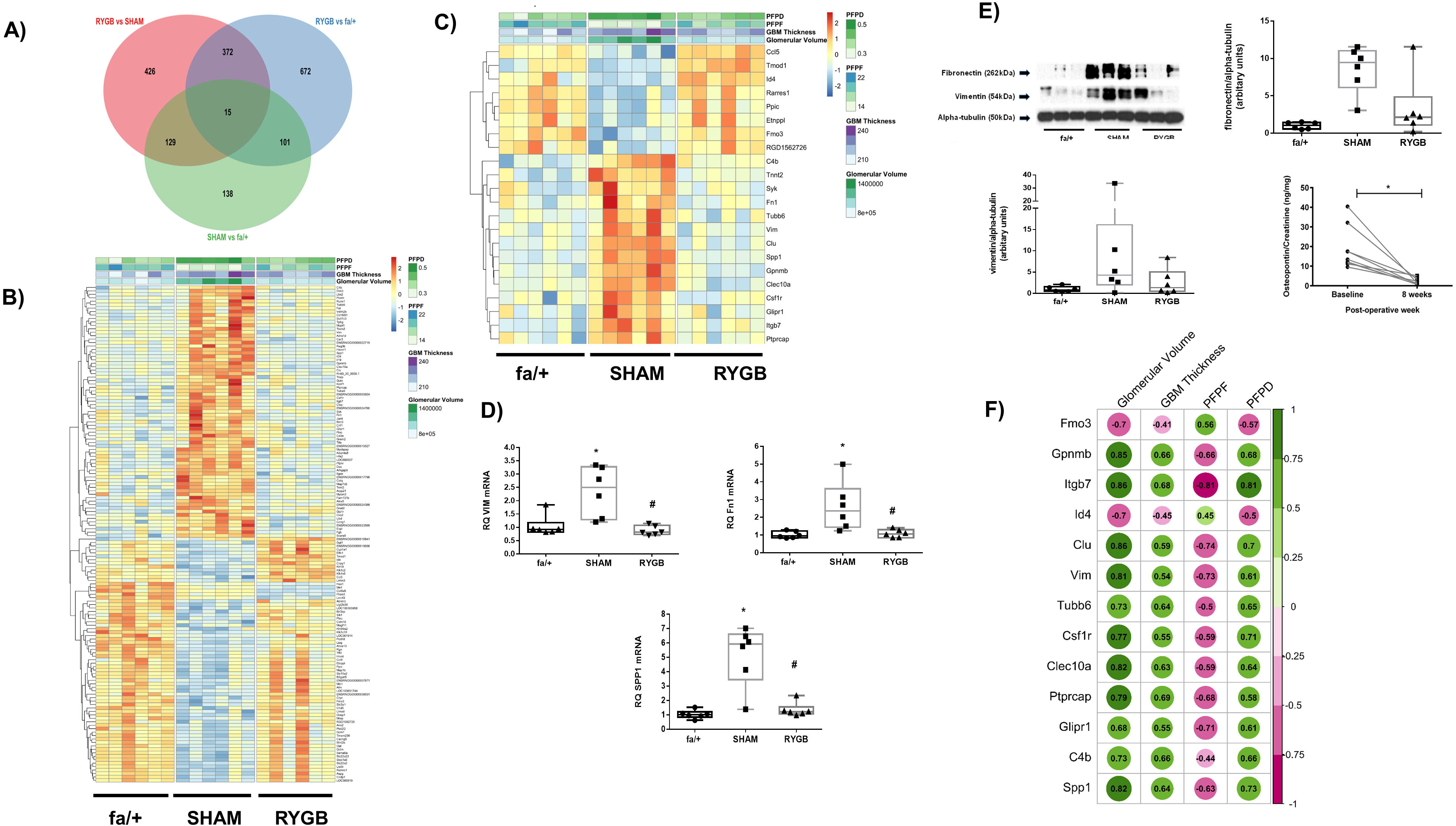
RYGB Normalises the Expression of Genes that Correlate with Markers of Renal Injury and Cross-reference to the Human DKD Transcriptome. A) Venn diagram showing number of gene changes in each differential expression analysis and extent of overlap. B) Heat-map of expression of 144 genes significantly changed in both SHAM versus fa/+ and RYGB versus SHAM comparisons. C) Heat-map of expression of genes changed in both comparisons that cross-reference to significantly altered expression profiles in microarray analysis of glomerular isolates from 13 healthy living donors, and 9 diabetic nephropathy patients. (Nephroseq-Woroniecka dataset). In both B and C, colour intensity for each sample in the heat-maps reflects relative abundance of given transcript based on normalised counts from RNA-Seq with blue indicating lower expression and orange-red indicating higher expression. Heat maps are headed by annotation of quantitative measures of glomerular structure and ultrastructure for each sample included in the analysis. Validation of expression changes in vimentin, fibronectin and Spp1 was undertaken at the mRNA level by qRT-PCR (D) and Western blotting (E). F) Correlation of expression levels for human validated genes from panel C with quantitative measures of glomerular injury in sections of rat renal cortex. Green indicates a positive correlation and purple a negative correlation. Increasing colour intensity and spot-size signifies increasing correlation coefficient. SHAM-sham surgery (laparotomy); RYGB-Roux-en-Y Gastric Bypass, *p<0.05 vs fa/+; #p<0.05 vs. SHAM.For osteopontin ELISA *p<0.05 baseline versus post-operative week 8.

### Suppression of TGF-β1 Target Gene Expression in the Kidney Following RYGB Coincides with Transcriptional Evidence of Restoration of BMP-7 Signalling (Figures 4)

Figure 5A presents a heatmap of z-scores of the top 10 upstream regulators of pathway enrichment for SHAM versus fa/+ and RYGB versus SHAM comparisons. In line with previous observations, and indicative of the primacy of changes in extracellular matrix pathway activation between fa/+, SHAM and RYGB groups, TGF-β1 signalling was identified as a key predicted upstream regulator in disease which is prominently suppressed following RYGB. Notably, RYGB also reversed the upstream regulator influence of several inflammation related factors including CSF2, IL-1β and TNFα. In addition, the c-myc related cell cycle regulating factor ZBTB17 predicted to act as an upstream regulator of transcriptional change in the kidney following RYGB, an observation coherent with the strong pathway level signal for changes in cell cycle pathway activation.

**Figure 5-.**
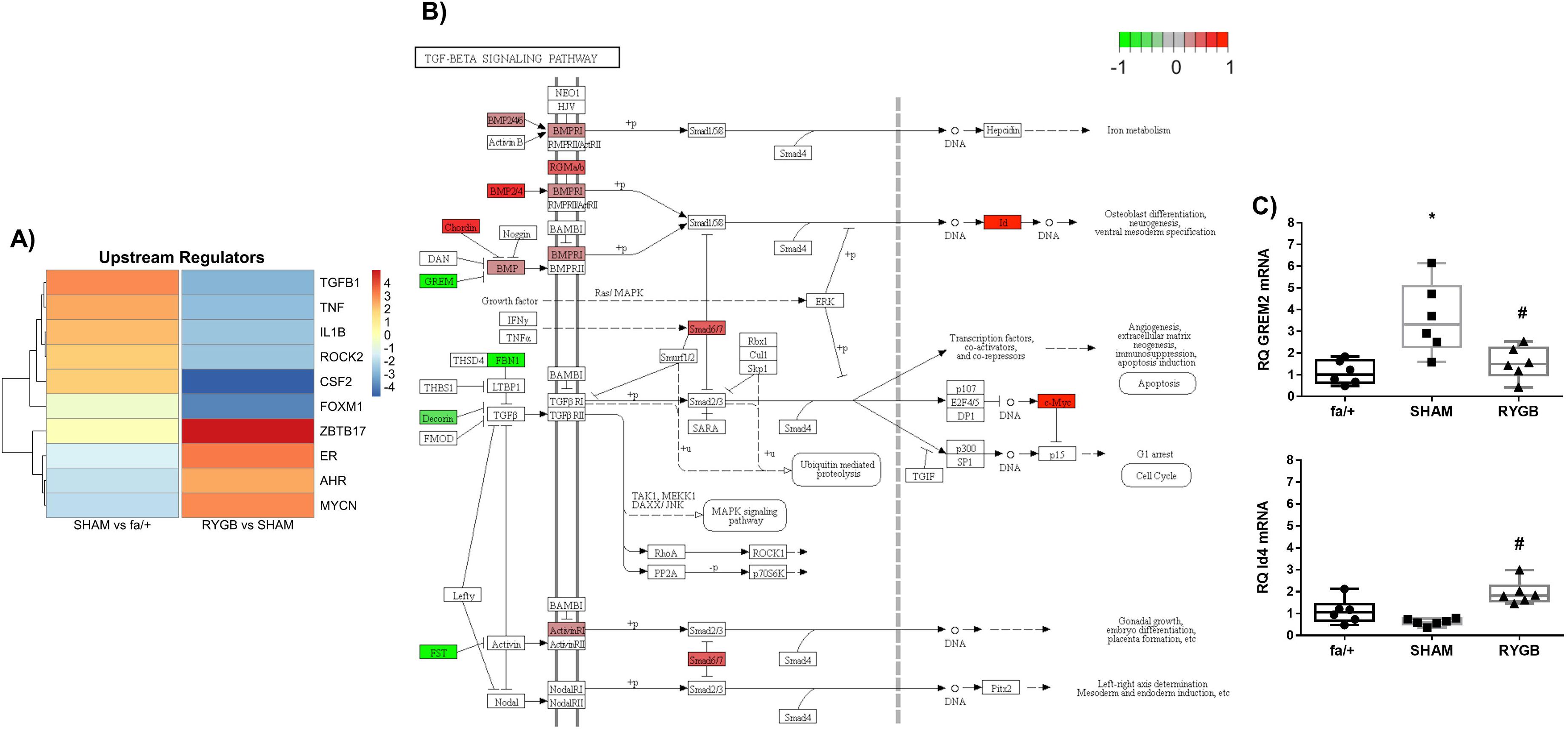
TGF-β1 Pathway activation is Reduced Following RYGB. A) The upstream regulator function of Ingenuity Pathway Analysis software was used to predict master transcriptional regulators underpinning disease associated shifts in the renal cortical transcriptome in SHAM operated rats and changes in the disease associated transcriptome arising secondary to RYGB. Colour intensity reflects weighted z-score. (B) Annotation of the KEGG TGF-β signalling pathway with genes, the expression of which is significantly altered by RYGB. Green signifies that expression is reduced by RYGB and red indicates increased expression following RYGB. (SHAM-sham surgery (laparotomy); RYGB-Roux-en-Y Gastric Bypass . C) Validation of expression changes in Grem-2 and Id4 at the mRNA level was undertaken by qRT-PCR.). *p<0.05 SHAM vs fa/+ and #p<0.05 RYGB vs. SHAM.

We visualised constituent changes in the KEGG TGF-β signalling pathway via Pathview for the RYGB versus SHAM comparison (Figure 5B). RYGB reduced expression of genes involved in the regulated sequestration and activation of TGF-β1 including fibrillin 1 (FBN1), Decorin and Follistatin. RYGB also reduced the expression of the Bone-morphogenic protein (BMP) antagonist Gremlin-2 and increased the expression of the BMP-7 responsive Id4 gene, suggestive of restoration of anti-fibrotic signalling in the renal tubule. Tandem reciprocal changes in Gremlin-2 and Id4 transcript levels following RYGB were validated by qPCR analysis (Figure 5C)

## DISCUSSION

We sought to extend insights into the clinically observed effects of RYGB in patients with DKD using a reverse translational research approach. We corroborated our previous findings regarding the beneficial impact of RYGB on the cardinal biochemical, structural and ultrastructural features of experimental DKD in the ZDF rat. (21, 22) We sought then to describe both the transcriptional landscape in the kidneys of ZDF SHAM rats, the changes to that landscape arising secondary to RYGB and how these changes correlated with structural and ultrastructural indicators of renal injury.

We identified 379 mRNA transcripts that were differentially regulated in the ZDF SHAM rat kidney versus fa/+ control. This number of genes is comparable in quantity and quality to previously reported changes at 28 weeks of age in whole renal tissue RNA-Seq from the related obese and diabetic Zucker/SHHF (ZS) rat (26); ZDF-167 genes up 212 genes down and ZS rats 184 genes up, 396 genes down. The predominance of proinflammatory and pro-fibrotic signals observed in ZDF SHAM animals and the implication of TGF-β1 signalling as an upstream regulator is also shared with the analysis from ZS rats (26). Whilst single cell RNA-Seq studies of murine and human diabetic kidney disease have recently been published (27, 28), the capacity for comparative extrapolation of our data to these profiles is limited, given the likely predominance of tubular epithelial derived transcripts in the renal parenchymal signal in our dataset. However, of the 379 differentially expressed genes in the ZDF SHAM rat kidney, 77 were significantly altered in the same direction as in the Woroniecka DKD bulk glomerular dataset, showing that transcriptional changes in the whole renal tissue of rats showed significant homology to transcriptional profiles in human DKD. (25)

We demonstrated that RYGB has an impact on the renal cortical transcriptome in DKD, correcting 38% of the disease associated changes in the transcriptome and reversing activation of many of the pathway level changes occurring in disease and their upstream regulators. In this respect, the pre-eminence of patterns of transcriptional change that map at the pathway level to processes governing inflammation, biological oxidation and, most notably, the fibrotic response is indicative of RYGB being associated with a broad range of renoprotective effects. Reduced activation of the TGF-β1 signalling pathway in the kidney after RYGB is a key finding and remphasises both the importance of this pathway in the progression of DKD, and involvement of interruption of this pathway as a common mechanism driving the anti-fibrotic effects of established therapies such as renin-angiotensin-aldosterone system blockade. (29)

Importantly, those RYGB-induced improvements in the renal transcriptome that translated to the human DKD transcriptome also correlated with improvements in biochemical, histological and ultrastructural markers of glomerular damage. This is interesting given that tubular injury related signals likely predominate in our dataset and supports the concept that although glomerular damage in early DKD is readily apparent at both the biochemical and histological level, these changes may be triggered by renal tubular dysfunction in early DKD through tubulo-glomerular cross-talk. (30) The specific responses of the renal tubule to RYGB should be the focus of future studies, particularly the impact of improved metabolic control on mitochondrial bioenergetics and the metabolic switch away from fatty acid oxidation that is now proposed to underpin renal tubular dedifferentiation and pro-fibrotic activation in DKD. (31).

Our data show coherent downward shifts in renal mRNA expression of the *Spp1* gene and the urinary excretion of its protein product osteopontin after RYGB, suggesting the potential utililty of this protein as a marker of treatment response. In addition, post-operative improvements in the mRNA expression of other cardinal indicators of tubular injury and recovery such as *Havcr1* (KIM-1, four fold reduction in RYGB versus Sham) and Umod (Uromodulin, two-fold increase in RYGB versus Sham) points to the potential utility of treatment responsive biomarker assessment which could be used in parallel with measurement of urinary albumin excretion as a non-invasive means of monitoring the short to medium term therapeutic response to surgery. Serum uromodulin has already been identified as a potentially useful marker of renal improvement post-metabolic surgery in patients with T2DM and obesity (32)

The comprehensive transcriptome wide profiling approach adopted herein have permitted us to that approximately 86% of changes to the renal transcriptome induced by RYGB are unique to the procedure itself and do not simply reflect reversal of disease-associated gene expression patterns. Some of the alterations in expression signatures particular to the RYGB intervention may be related to unique beneficial effects of the procedure, and interrogation of candidates in this regard will form the basis of future mechanistic studies. Conversely, a proportion of these changes likely reflect adaptive responses to undesirable effects of the procedure. Owing to the anatomical reconfiguration occurring in RYGB, uptake of certain nutrients and minerals including iron, calcium and vitamin D which are normally absorbed in the proximal small intestine can be impaired. (33–35) For example, calcium absorption efficacy in the proximal jejunum is regulated by activated vitamin D via the vitamin D receptor. (36) RNAseq analysis revealed that one of the top differentially expressed genes in the RYGB ZDF rat kidney was Cyp27b1. Cyp27b1 encodes 25-Hydroxyvitamin D3 1-α-hydroxylase, the enzyme that converts vitamin D to its active form 1,25-dihydroxyvitamin D3. Upregulated expression of the enzyme in the RYGB kidney suggests that there could be resistance to the effects of 1,25 dihydroxy-vitamin D after RYGB, which thus prevents normal negative feedback inhibition of Cyp27b1 in the vitamin D replete state. (37) Future studies directed at differentiating the beneficial from the detrimental effects of RYGB on the kidney via interrogation of the RYGB-specific transcriptional changes and assessment of their likely impact on the kidney using *in vitro* modelling merit consideration.

There are some noteworthy limitations of our study. As a result of loss of leptin deficiency and attendant reductions in sympathetic drive, the ZDF rat is not hypertensive until much later in the process of renal decline and thus our data are likely more relevant to normotensive or blood pressure controlled DKD. That said, RYGB effectively treats hypertension (38), a phenomenon that may be attributable in part to enhanced natriuresis (19–20). The ZDF rat also does not display prominent tubulointerstitial fibrosis and renal functional decline until late in the course of the disease. However, pathway analysis of the correction in the renal transcriptome induced by RYGB highlights interruption of the early pro-fibrotic programme as a key effect of the intervention. Definitive resolution of the effect of RYGB on discrete cellular subsets in the diabetic kidney was not possible in the present study but future studies using single cell RNA-Seq are warranted.

This study defines key transcriptomic and pathway level changes that characterise the arrest of DKD progression following RYGB. In future it may be possible to replicate these transcriptomic and pathway level changes without the use of surgery. An important intermediate step in attempting to achieve this would involve generating a comprehensive assessment of the activation state of implicated signalling pathways involved, ideally using an unsupervised phospho-proteomics bioinformatic analysis based approach which may allow for proximate novel receptor level targets to be identified for rational drug design. The current mechanistic evidence however supports the potential of RYGB to form a part of the treatment algorithm targeted at addressing the burden of renal disease arising from increases in the prevalence of obesity and type 2 diabetes.

## Supporting information

Supplementary File 1- Bioinformatic and Pathway Enrichment Analyses.

## ACKNOWLEDGEMENTS

NGD, LF and CIR devised and designed the studies. MN, JE, NF, HGE and NGD performed surgery and animal husbandry. MN conducted biochemical assays, RNA-Seq, histological and immunohistochemical studies with support from HE, JMcC and WPM. Bioinformatic pathway and upstream regulator analyses were conducted by WPM, VZ and EPB. Validation of transcriptomic signals at mRNA and protein level was conducted by MN and WPM. NGD, MN, WPM, EPB, LF, CG and CIR analysed and interpreted data. NGD, WPM, MN and CIR drafted the manuscript with critical input from CG, EPB and LF.

We acknowledge the expert guidance and technical support to MN and WM with regards to Western blotting received from Ms Mariam Marai (University College Dublin) and Associate Professor Anna Casselbrant (University of Gothenburg). We also express our gratitude for the expert support of Dr. Sanna Abrahamsson from the Bioinformatics Core Facility at the Sahlgrenska Academy, University of Gothenburg, Sweden who handled RNA-Seq raw data processing and differential expression analyses. We also acknowledge local support received in the realisation of these studies from all at the UCD Biomedical Facility and Dr.’s Catherine Moss and Alison Murphy at the Genomics Core Facility at The UCD Conway Institute.

Funding support from the following agencies is acknowledged; Science Foundation Ireland (12/YI/B2480) to CIR, Swedish Medical Research Council (2015-02733) and European Foundation for the Study of Diabetes /Boehringer Ingelheim European Diabetes Research Programme (BI 2017_3) to CIR and NGD, Science Foundation Ireland (15/IA/3152 and 15/US/B3130) to CG and EPB. WPM’s contribution was performed within the Irish Clinical Academic Training (ICAT) Programme, supported by the Wellcome Trust and the Health Research Board (Grant Number 203930/B/16/Z), the Health Service Executive National Doctors Training and Planning and the Health and Social Care, Research and Development Division, Northern Ireland.

NGD and CIR are co-guarantors of this work and, as such, had full access to all the data in the study and take responsibility for the integrity of the data and the accuracy of the data analysis.

CIR discloses personal fees outside of the submitted work from Novo Nordisk, Gl Dynamics, Eli Lilly, Johnson and Johnson, Sanofi, Aventis, Astra Zeneca, Janssen, Bristol-Myers Squibb and Boehringer-Ingelheim

## Ethics Approval Disclosure Statement

All studies detailed were approved by the University College Dublin Animal Research Ethics Committee and licensed by Government of Ireland Health Products Regulatory Agency (protocol #AE18692-P084)

---

**Supplementary File 1: Bioinformatic and Pathway Enrichment Analyses.**

**Supplementary File 2:List of Differentially Expressed Genes (SHAM versus fa/+)**

**Supplementary File 3:List of Differentially Expressed Genes (RYGB versus SHAM)**

## Notes

https://www.ncbi.nlm.nih.gov/geo/query/acc.cgi?acc=GSE117380

### Competing Interest Statement

ClR discloses personal fees outside of the submitted work from Novo Nordisk, GI Dynamics, Eli Lilly, Johnson and Johnson, Sanofi, Aventis, Astra Zeneca, Janssen, Bristol-Myers Squibb and Boehringer-Ingelheim

## REFERENCES

1. Global, regional, and national burden of chronic kidney disease, 1990-2017: a systematic analysis for the Global Burden of Disease Study 2017. Lancet 2020;395:709–733

2. Saran R, Robinson B, Abbott KC, Agodoa LYC, Bragg-Gresham J, Balkrishnan R, et al. US Renal Data System 2018 Annual Data Report: Epidemiology of Kidney Disease in the United States. American journal of kidney diseases : the official journal of the National Kidney Foundation. 2019;73(3s1):A7–a8.

3. Afkarian M, Sachs MC, Kestenbaum B, Hirsch IB, Tuttle KR, Himmelfarb J, et al. Kidney disease and increased mortality risk in type 2 diabetes. Journal of the American Society of Nephrology : JASN. 2013;24(2):302–8.

4. Stratton IM, Cull CA, Adler AI, Matthews DR, Neil HA, Holman RR. Additive effects of glycaemia and blood pressure exposure on risk of complications in type 2 diabetes: a prospective observational study (UKPDS 75). Diabetologia. 2006;49(8):1761–9.

5. Oellgaard J, Gaede P, Rossing P, Persson F, Parving HH, Pedersen O. Intensified multifactorial intervention in type 2 diabetics with microalbuminuria leads to long-term renal benefits. Kidney international. 2017;91(4):982–8.

6. Perkovic V, Jardine MJ, Neal B, Bompoint S, Heerspink HJL, Charytan DM, et al. Canagliflozin and Renal Outcomes in Type 2 Diabetes and Nephropathy. N Engl J Med. 2019;380(24):2295–306.

7. Sarafidis PA, Tsapas A. Empagliflozin, Cardiovascular Outcomes, and Mortality in Type 2 Diabetes. N Engl J Med. 2016;374(11):1092.

8. Mosenzon O, Wiviott SD, Cahn A, Rozenberg A, Yanuv I, Goodrich EL, et al. Effects of dapagliflozin on development and progression of kidney disease in patients with type 2 diabetes: an analysis from the DECLARE-TIMI 58 randomised trial. The lancet Diabetes & endocrinology. 2019;7(8):606–I7.

9. Mann JFE, Orsted DD, Brown-Frandsen K, Marso SP, Poulter NR, Rasmussen S, et al. Liraglutide and Renal Outcomes in Type 2 Diabetes. N Engl J Med. 2017;377(9):839–48.

10. Ikramuddin S, Korner J, Lee WJ, Connett JE, Inabnet WB, Billington CJ, et al. Roux-en-Y gastric bypass vs intensive medical management for the control of type 2 diabetes, hypertension, and hyperlipidemia: the Diabetes Surgery Study randomized clinical trial. JAMA. 2013;309(21):2240–9.

11. Mingrone G, Panunzi S, De Gaetano A, Guidone C, laconelli A, Leccesi L, et al. Bariatric surgery versus conventional medical therapy for type 2 diabetes. N Engl J Med. 2012;366(17):1577–85.

12. Schauer PR, Bhatt DL, Kirwan JP, Wolski K, Aminian A, Brethauer SA, et al. Bariatric Surgery versus Intensive Medical Therapy for Diabetes - 5-Year Outcomes. N Engl J Med. 2017;376(7):641–51.

13. Rubino F, Nathan DM, Eckel RH, Schauer PR, Alberti KG, Zimmet PZ, et al. Metabolic Surgery in the Treatment Algorithm for Type 2 Diabetes: a Joint Statement by International Diabetes Organizations. Obes Surg. 2017;27(1):2–21.

14. Martin WP, Docherty NG, Le Roux CW. Impact of bariatric surgery on cardiovascular and renal complications of diabetes: a focus on clinical outcomes and putative mechanisms. Expert review of endocrinology & metabolism. 2018;13(5):251–62.

15. Billeter AT, Scheurlen KM, Probst P, Eichel S, Nickel F, Kopf S, et al. Meta-analysis of metabolic surgery versus medical treatment for microvascular complications in patients with type 2 diabetes mellitus. The British Journal of Surgery 2018; 105:168–181

16. Upala S, Wijarnpreecha K, Congrete S, Rattanawong P, Sanguankeo, A. Bariatric surgery reduces urinary albumin excretion in diabetic nephropathy: a systematic review and meta-analysis. Surgery for obesity and related diseases : official journal of the American Society for Bariatric Surgery. 2016; 12:1037–1044,

17. Bjornstad P, Hughan K, Kelsey MM, Shah AS, Lynch J, Nehus E, et al. Effect of surgical versus medical therapy on diabetic kidney disease over 5 Years in severely obese adolescents with type 2 diabetes. Diabetes Care 2020; 43:187–195

18. Cohen RV, Veiga Pereira T, Mamedio Aboud C, Zanata Petry TB, Lopes Correa JL, Schiavo CA, et al. Effect of Gastric Bypass vs Best Medical Treatment on Early-Stage Chronic Kidney Disease in Patients With Type 2 Diabetes and Obesity A Randomized Clinical Trial. JAMA Surgery 2020; in press, doi:doi:10.1001/jamasurg.2020.0420

19. Docherty NG, Fandriks L, le Roux CW, Hallersund P, Werling M. Urinary sodium excretion after gastric bypass surgery. Surgery for obesity and related diseases : official journal of the American Society for Bariatric Surgery. 2017;13(9):1506–14.

20. Hallersund P, Sjostrom L, Olbers T, Lonroth H, Jacobson P, Wallenius V, et al. Gastric bypass surgery is followed by lowered blood pressure and increased diuresis - long term results from the Swedish Obese Subjects (SOS) study. PloS one. 2012;7(11):e49696.

21. Canney AL, Cohen RV, Elliott JA, C MA, Martin WP, Docherty NG, et al. Improvements in diabetic albuminuria and podocyte differentiation following Roux-en-Y gastric bypass surgery. Diabetes & vascular disease research. 2019:1479164119879039.

22. Neff KJ, Elliott JA, Corteville C, Abegg K, Boza C, Lutz TA, et al. Effect of Roux-en-Y gastric bypass and diet-induced weight loss on diabetic kidney disease in the Zucker diabetic fatty rat. Surgery for obesity and related diseases : official journal of the American Society for Bariatric Surgery. 2017;13(1):21–7.

23. Lane PH, Steffes MW, Mauer SM. Estimation of glomerular volume: a comparison of four methods. Kidney international. 1992;41(4):1085–9.

24. Haas M. Thin glomerular basement membrane nephropathy: incidence in 3471 consecutive renal biopsies examined by electron microscopy. Archives of pathology & laboratory medicine. 2006;130(5):699–706.

25. Woroniecka KI, Park ASD, Mohtat D, Thomas DB, Pullman JM, Susztak K. Transcriptome Analysis of Human Diabetic Kidney Disease. Diabetes. 2011;60(9):2354–69.

26. Kelly KJ, Liu Y, Zhang J, Goswami C, Lin H, Dominguez JH. Comprehensive genomic profiling in diabetic nephropathy reveals the predominance of proinflammatory pathways. Physiological genomics. 2013;45(16):710–9.

27. Wilson PC, Wu H, Kirita Y, Uchimura K, Ledru N, Rennke HG, et al. The single-cell transcriptomic landscape of early human diabetic nephropathy. Proceedings of the National Academy of Sciences of the United States of America. 2019;116(39):19619–25.

28. Fu J, Akat KM, Sun Z, Zhang W, Schlondorff D, Liu Z, et al. Single-Cell RNA Profiling of Glomerular Cells Shows Dynamic Changes in Experimental Diabetic Kidney Disease. Journal of the American Society of Nephrology : JASN. 2019;30(4):533–45.

29. Docherty NG, Murphy M, Martin F, Brennan EP, Godson C: Targeting cellular drivers and counter-regulators of hyperglycaemia- and transforming growth factor-beta1-associated profibrotic responses in diabetic kidney disease. Experimental physiology 2014;99:1154–1162

30. Zeni L, Norden AGW, Cancarini G, Unwin RJ. A more tubulocentric view of diabetic kidney disease. Journal of nephrology. 2017;30(6):701–I7.

31. Kang HM, Ahn SH, Choi P, Ko YA, Han SH, Chinga F, et al. Defective fatty acid oxidation in renal tubular epithelial cells has a key role in kidney fibrosis development. Nature medicine. 2015;21(1):37–46.

32. Scheurlen KM, Billeter AT, Kopf S, Herbst V, Block M, Nawroth PP, Zeier M, Scherberich JE, Müller-Stich BP: Serum uromodulin and Roux-en-Y gastric bypass: improvement of a marker reflecting nephron mass. Surgery for obesity and related diseases : official journal of the American Society for Bariatric Surgery 2019;15:1319–1325

33. Xanthakos SA. Nutritional Deficiencies in Obesity and After Bariatric Surgery. Pediatric clinics of North America. 2009;56(5):1105–21.

34. Decker GA, Swain JM, Crowell MD, Scolapio JS. Gastrointestinal and nutritional complications after bariatric surgery. Am J Gastroenterol. 2007;102(11):2571-80; quiz 81.

35. Gesquiere I, Lannoo M, Augustijns P, Matthys C, Van der Schueren B, Foulon V. Iron deficiency after Roux-en-Y gastric bypass: insufficient iron absorption from oral iron supplements. Obes Surg. 2014;24(1):56–61.

36. Elias E, Casselbrant A, Werling M, Abegg K, Vincent RP, Alaghband-Zadeh J, et al. Bone mineral density and expression of vitamin D receptor-dependent calcium uptake mechanisms in the proximal small intestine after bariatric surgery. The British journal of surgery. 2014;101(12):1566–75.

37. Murayama A, Takeyama K, Kitanaka S, Kodera Y, Kawaguchi Y, Hosoya T, et al. Positive and negative regulations of the renal 25-hydroxyvitamin D3 1alpha-hydroxylase gene by parathyroid hormone, calcitonin, and 1alpha,25(OH)2D3 in intact animals. Endocrinology. 1999;140(5):2224–31.

38. Schiavon CA, Bersch-Ferreira AC, Santucci EV, Oliveira JD, Torreglosa CR, Bueno PT, et al. Effects of Bariatric Surgery in Obese Patients With Hypertension: The GATEWAY Randomized Trial (Gastric Bypass to Treat Obese Patients With Steady Hypertension). Circulation. 2017.

