## Supplementary File 1- Bioinformatic and Pathway Enrichment Analyses. for "Characterisation of the Renal Cortical Transcriptome Following Roux-en-Y Gastric Bypass Surgery in Experimental Diabetic Kidney Disease"

The quality of the reads were examined using fastqc/0.11.2 (http://www.bioinformatics.babraham.ac.uk/projects/fastqc), the resulting reports were merged using multiqc/0.9 [1]. The reads were quality filtered using prinseq/0.20.3 [2]. The quality filtered reads were mapped towards the rat reference genome (rn5) with STAR/2.5.2b [3]. HTseq/0.6.1p [4] was used for quantifying the amount of reads mapped towards the genes.

Further analysis was performed using the R statistical programming language. Differential expression analysis was performed in DESeq2 [5]. The data was normalized using sizefactors. A generalized linear model was fitted to the normalized data. Wald statistics was used as the statistical test. The p-values were adjusted for multiple testing with the Benjamini-Hochberg procedure.

Principal component analysis was performed using DESeq2-normalized gene expression counts data. Volcano plots were generate using the R package EnhancedVolcano [6]. Differentially expressed transcripts with absolute fold-change ≥1.3 and adjusted p-value <0.05 were selected for pathway enrichment analysis using Reactome and KEGG databases via the R packages ReactomePA and clusterProfiler [7,8]. Reactome pathway enrichment results are presented on dotplots ranking pathways by statistical significance, number of pathway-specific transcripts changed, and gene ratio. Enriched Reactome pathways were compared between SHAM vs fa/+ and RYGB vs SHAM differentially expressed genes using clusterProfiler. KEGG pathways enriched in our dataset were visualsied using pathview [9]. Upstream regulator analysis was performed using Ingenuity Pathway Analysis software. Rat transcript IDs were converted to their human orthologs to estimate the abundance of renal cortical immune and stromal cell populations using MCPcounter [10].

Differentially expressed transcripts (absolute fold-change ≥1.3, adjusted p-value <0.05) between study groups were visualized on Venn diagrams. Transcripts which changed from health to disease (SHAM vs fa/+) and again with RYGB surgery (RYGB vs SHAM) were selected. A publicly available list of differentially expressed genes (absolute fold-change ≥1.5, adjusted p-value <0.05) in glomeruli of people with DKD versus healthy controls (the Woroniecka human DKD glomerular microarray dataset) were converted to their rat orthologs and intersected with genes overlapping between SHAM vs fa/+ and RYGB vs SHAM comparisons [11]. Expression of these genes in our dataset was plotted using pheatmap [12]. The relationship between gene expression and glomerular structure was interrogated using Pearson correlations by creating a correlation matrix of DESeq2-normalized gene expression counts and numerical data from histology/TEM. Pearson r values were plotted using corrplot [13].

1. Ewels, P., et al., *MultiQC: Summarize analysis results for multiple tools and samples in a single	report*. Bioinformatics, 2016, 32(19): p. 3047–3048
2. Schmieder, R. and R. Edwards, *Quality control and preprocessing of metagenomic datasets.* Bioinformatics, 2011. **27**(6): p. 863-4.
3. Dobin, A., et al., *STAR: ultrafast universal RNA-seq aligner.* Bioinformatics, 2013. **29**(1): p. 15-21.
4. Anders, S., P.T. Pyl, and W. Huber, *HTSeq--a Python framework to work with high-throughput sequencing data.* Bioinformatics, 2015. **31**(2): p. 166-9.
5. Love, M.I., W. Huber, and S. Anders, *Moderated estimation of fold change and dispersion for RNA-seq data with DESeq2.* Genome Biol, 2014. **15**(12): p. 550.
6. Kevin Blighe (2019). EnhancedVolcano: Publication-ready volcano plots with enhanced

colouring and labeling. R package version 1.2.0. <https://github.com/kevinblighe/EnhancedVolcano>

7. Guangchuang Yu, Qing-Yu He. ReactomePA: an R/Bioconductor package for reactome

pathway analysis and visualization. Molecular BioSystems 2016, 12(2):477-479.

8. Guangchuang Yu, Li-Gen Wang, Yanyan Han and Qing-Yu He. clusterProfiler: an R package for comparing biological themes among gene clusters. OMICS: A Journal of Integrative Biology 2012, 16(5):284-287.
9. Luo, W. and Brouwer C., Pathview: an R/Bioconductor package for pathway-based data

integration and visualization. Bioinformatics, 2013, 29(14): 1830-1831, doi:10.1093/bioinformatics/btt285

10. Etienne Becht and Aurelien de Reynies (2016). MCPcounter: Estimating

tissue-infiltrating immune and other stromal subpopulations abundances using gene expression. R package version 1.1.0.

11. Woroniecka KI, Park ASD, Mohtat D, Thomas DB, Pullman JM, Susztak K. Transcriptome Analysis of Human Diabetic Kidney Disease. Diabetes. 2011;60(9):2354-69.
12. Raivo Kolde (2019). pheatmap: Pretty Heatmaps. R package version 1.0.12. <https://CRAN.R-project.org/package=pheatmap>.
13. Taiyun Wei and Viliam Simko (2017). R package "corrplot": Visualization of a

Correlation Matrix (Version 0.84). Available from https://github.com/taiyun/corrplot.
